# VE-cadherin NOT-gated CD93 CAR T cells discriminate between AML and healthy endothelial cells

**DOI:** 10.64898/2026.03.09.710630

**Authors:** Tess Woodring, Otto A. Kletzien, Kierstin Schlevensky, Cristina Sanchez-de-Diego, Paresh Vishwasrao, Lauren Mahoney, Sheena C Kerr, Rebecca M. Richards

## Abstract

**Background:** Chimeric antigen receptor (CAR) T cell therapy has transformed the treatment of B cell malignancies, but translation to acute myeloid leukemia (AML) has been hindered by on-target, off-tumor (OTOT) toxicity. In particular, endothelial cell (EC)-specific toxicity has limited clinical translation of promising leukemia stem cell-enriched targets such as CD93. Innovative strategies to mitigate EC damage while preserving antileukemic efficacy are needed.

**Methods:** We hypothesized that a NOT-gated CAR T cell strategy could circumvent EC toxicity associated with CD93 targeting. Considering CAR target antigen density and the pro-inflammatory microenvironment of CAR T cells, we identified VE-cadherin (VC), a highly specific EC marker, as an optimal inhibitory CAR target. We engineered a novel VC-specific single chain variable fragment (scFv), confirmed EC specificity in the context of a VC-specific second-generation activating CAR, then evaluated VC/CD93 NOT-gated CAR T cells for EC protection and antileukemic activity in *in vitro* cytotoxicity assays and in a three-dimensional vascularized microphysiological system.

**Results:** VC/CD93 NOT-gated CAR T cells maintain potent cytotoxicity against AML across multiple effector-to-target ratios, but preserve EC integrity, including in a three-dimensional vascular model system. Importantly, prior AML exposure did not impair the EC-protective function of the VC-specific iCAR, indicating durable NOT-gate activity under inflammatory conditions. Conversely, EC-induced iCAR inhibitory functions did not limit downstream antileukemic cytotoxicity, confirming a reversibility of both activation and inhibitory signals. *Conclusions:* These findings establish NOT-gated CAR T cells as an effective strategy to overcome EC-specific OTOT toxicity. Our results underscore the importance of CAR target discovery and validation across a spectrum of inflammatory states that can influence antigen expression and available therapeutic windows. This approach expands the potential CAR target landscape for AML and may be more broadly applicable to other malignancies where OTOT toxicity limits clinical translation.

## Background

Most patients with acute myeloid leukemia (AML) will die within 5 years of diagnosis (1, 2). Molecular therapies like FLT inhibitors (e.g. gilteritinib) and pro-apoptotic agents (e.g. venetoclax) have improved overall response rate and survival with first-line therapy but have yet to close the gap in outcomes between AML and other acute leukemias like B-cell acute lymphoblastic leukemia (B-ALL) (3). While allogeneic hematopoietic stem cell transplant (HSCT) has advanced significantly over the past decades, providing both intensive cytoreduction and graft-versus-tumor effect, there remains no effective salvage therapy for patients with AML who relapse after HSCT (4).

Chimeric antigen receptor (CAR) T cells are a promising option for meeting the urgent need for new treatments. CAR T cells couple high-affinity antigen-recognition with T cell cytotoxicity, providing targeted cancer killing that bypasses the need for antigen presentation via the major histocompatibility complex (MHC). Their efficacy in aggressive B cell-derived blood cancers has led to unprecedented cures and seven FDA-approved CAR T cell therapies as of 2025 (5, 6). Nonetheless, even the most successful CAR T cells exert on-target, off-tumor (OTOT) toxicity. In the case of CD19 CAR T cells, OTOT toxicity affects healthy B cells, the clinical consequences of which can be supported with immunoglobulin replacement. In contrast, most AML targets have overlapping expression patterns with more critical healthy tissues, such as hematopoietic stem cells or endothelial cells (7, 8). Increasingly sophisticated CAR T cell engineering strategies to avoid OTOT toxicity provide avenues to further develop targets previously deemed too high risk, thus expanding treatment opportunities for AML.

We have previously shown CD93 to be an appealing CAR T-cell target expressed on AML blasts and leukemic stem cells (LSCs) but not hematopoietic stem and progenitor cells (HSPCs), offering leukemic killing with potential to avoid prolonged marrow suppression (9). However, CD93 is also expressed on healthy human endothelial cells (ECs), which poses a direct risk for unacceptable vascular damage. One strategy for circumventing this problem is Boolean logic-based NOT-gating, in which a CD93-specific CAR activation signal is combined with an opposing EC-specific inhibitory signal (*a* NOT *b*) to prevent OTOT activation. The NOT-gating approach can enhance specificity without narrowing the breadth of on-target tumor cell killing or adding new opportunities for tumor antigen escape, as would occur with alternative logic gating strategies such as AND-gating. On a mechanistic level, NOT-gating is achieved by introduction of inhibitory CARs (iCARs). These chimeric receptors leverage immune cell-derived inhibitory signaling after healthy-cell target recognition in a process analogous to activation of intrinsic T-cell activation domains by activating CARs (aCARs) upon tumor engagement (10–13). Using CD19 as a surrogate endothelial-specific target, we previously designed NOT-gated CAR T cells that could abrogate CD93 CAR T cell activity against ECs with a CD19-specific iCAR (9). However, the optimal endogenous EC-specific ligand to engage the iCAR remained unknown, and the ability of NOT-gated AML CAR T cells to exhibit selective and regulatable cytotoxicity in a physiologically relevant model system had not been shown.

Here, we identified VE-cadherin (VC) as an ideal EC-specific ligand based on its differential expression in AML and ECs, both at rest and under pro-inflammatory conditions. We engineered VC/CD93 NOT-gated CAR T cells that kill AML cells and spare ECs as predicted by transcriptional and protein-level antigen expression. In contrast to other iCAR constructs that suppressed OTOT cytokine production but not cytotoxicity (14), our VC iCAR blunted both cytokine production and cytotoxicity upon exposure to ECs, while leaving intact the response to AML target cells. Importantly, VC iCARs were able to protect ECs even after killing AML target cells, and kill AML after bypassing EC damage. These results open a path to clinical translation of CD93 as a viable AML CAR target. Furthermore, our bioinformatics-guided target identification strategy is generalizable to the design of CAR T cells for other cancers in which tissue-specific OTOT toxicity limits access to potentially curative new immunotherapies.

## Methods

### Bulk RNASeq Analysis

Data from bulk RNASeq analysis was generated from two EC lines (iHUVEC, TIME) and three AML cell lines (Kasumi-1, NOMO-1, THP-1), as described previously (9). The sequencing dataset had been deposited in NCBI Gene Expression Omnibus (GEO) and accessible with the accession number GSE159991. Differential gene expression analysis comparing ECs and AML cells in each of the experimental conditions (unstimulated or “resting,” with IFNγ, with TNFα, and with both IFNγ and TNFα) was performed using DESeq2 within the GEO2R tool (15). Differentially expressed genes were filtered in R (version 4.4.2) using a statistical filter of false discovery rate (FDR) <0.05 and log2fold change >10. Genes were given molecular annotations as curated within the Human Surfacesome from the Wollschied laboratory (16). Gene expression levels were visualized using the ggplot2 package in R, and Venn diagrams were generated using the ggVennDiagram package in R.

### Healthy Gene Expression Atlas

Single-cell gene expression data from healthy cells was queried using the CZ CELLxGENE Discover tools for the complete atlas and endothelial cell compartments (17, 18).

### Patient AML RNASeq Data

Bulk RNASeq data for 3,225 primary patient AML samples in the TARGET-AML project (dbGaP Study Accession phs000218) was downloaded from the National Cancer Institute GDC Data Portal (19). Gene expression data (transcripts per million) was visualized for iCAR ligand candidates and AML markers (*CD33, FLT3, CLEC12A, CSF1R, CD86*) using ggplot2.

### Cell Lines

TIME cells were purchased from ATCC. AML cell lines were a gift from the labs of Ravi Majeti and Crystal Mackall (Stanford University). iHUVECs were gifted by Dr. David Sullivan at Northwestern University (Chicago, IL). HUVEC cells used in LumeNEXT experiments were purchased from Lonza. TIME and iHUVEC cell lines were stably transduced with a lentiviral vector containing mKate fused to a nuclear localization signal, followed by puromycin selection and flow cytometry-based sorting on an MA900 cell sorter (Sony). All cell lines underwent identity verification by short tandem repeat analysis and were confirmed to be *Mycoplasma* negative by PCR at least every 6 months.

### Flow Cytometry

All samples were analyzed with an Attune NxT flow cytometer (Thermo Fisher), and data were analyzed with FlowJo software. Fc block was applied prior to antibody staining for all AML and endothelial cell lines (BD Pharmingen). VE-cadherin was detected using CD144-APC (clone BV9, BioLegend), VEGFR2 was detected using CD309-PE (Clone RUO, Biolegend), and CD93 was detected using CD93-PE (Clone VIMD2, BioLegend). CD93 CAR was detected using His-tagged CD93 protein (Sino Biological) followed by APC-His antibody (Clone J095G46, BioLegend). VE-cadherin CAR was detected using VE-cadherin protein (Sino Biological) esterified to Dylight-488 (Thermo Fisher). Total CAR positivity was also measured using Alexa 647-Goat anti-human IgG Fab fragment antibody (Polyclonal, Jackson Immunoresearch). T-cell phenotype was evaluated using the following antibodies: CD4-BV711 (Clone OKT4, Biolegend), CD8-PE-Cy7 (Clone SK1, BD Biosciences), CD3-FITC (Clone HIT3a, Biolegend), PD-1-BV605 (Clone NAT105, Biolegend), LAG-3-PE (Clone 11C3C65, Biolegend), TIM-3-BV421 (Clone F38-2E2, Biolegend). Antigen density of CD93, VE-cadherin and VEGFR2 were measured by flow cytometry with the corresponding antibodies run alongside Quantibrite™ PE or APC quantitation beads (BD Biosciences), followed by normalization to the corresponding standard curves.

### Generation of CAR Constructs

The VE-cadherin (VC) antibody sequence from which the scFvs were derived was generously provided by Dr. Barry Gumbiner (University of Virginia), corresponding to the 2E11d mAb reported previously (20). The scFvs were designed by codon optimizing light and heavy chain sequences, and connecting them in either light-heavy or heavy-light orientation with a (G_4_S)_3_ linker sequence. Synthesis was performed by GeneArt (Invitrogen). The scFv DNA product was cloned into an MSGV1 retroviral vector with CD28 hinge, transmembrane, and intracellular domains along with the CD3ζ activating domain as previously described (21) to generate VC.28ζ. To generate the PD1 NOT-gated CAR (VC.PD1/CD93.28ζ), the VC scFv was inserted into the first position of a bicistronic retroviral vector containing CD8 hinge/transmembrane domains and the PD-1 inhibitory intracellular domain, followed by a P2A sequence. The second position of the bicistronic vector consisted of a CD93 scFv with CD28 hinge, transmembrane and intracellular domains adjacent to the CD3ζ activating domain. The Pdel control CAR (VC.Pdel/CD93.28ζ) was made by removing the PD-1 inhibitory intracellular domain from VC.PD1/CD93.28ζ). Each plasmid was transformed into One Shot™ Stbl3™ E. coli cells (Invitrogen). Colonies were picked and grown in 80 mL volumes for DNA purification by midiprep (Qiagen).

### CAR T-cell Transduction

Retroviral particles were generated by transfecting a HEK293 GP producer cell line (CVCL_E072) with both RD114 plasmid and the corresponding CAR plasmid. Viral supernatant was collected at 48 and 72 hours after transfection with fresh media added between the two collections. Cryopreserved primary human T cells had been obtained from a total of 9 healthy volunteers, purchased from Versiti (Milwaukee, WI) with consent (9). Human T cells were grown in AIM-V media supplemented with 100 IU/mL IL-2 and stimulated with CD3/CD28 Dynabeads™ (Gibco). 12-well non-coated plates were coated with Retronectin (Takara) at 4°C overnight, blocked with 2% BSA and treated with viral supernatant by centrifuging at 3200g at 32°C for 2-3 hours. Viral supernatant was removed and 0.25 to 0.5 million T cells were added to each well and cultured at 37°C overnight. The following day, a second round of transduction was performed, then CD3/CD28 beads were removed the following day by magnetic separation, and CARs were left to expand in fresh AIM-V media with IL-2 until characterization or functional assays performed between day 9 and 11 after activation.

### IncuCyte Cytotoxicity Assays

10,000 target cells (GFP-expressing AML cell lines or mKate-expressing endothelial cell lines) were co-cultured with CAR-T cells in triplicate wells at effector-to-target (E:T) ratios ranging from 1:2 to 1:4 in 150 μL media. AML co-cultures were grown in RPMI media supplemented with 10% FBS, 10 mM HEPES, 100 U/mL penicillin and 100 µg/mL streptomycin, while EC co-cultures were grown in a mix of 66% RPMI and 33% Vascular Cell Basal Media (ATCC) supplemented with the Microvascular Endothelial Growth Kit-VEGF (ATCC), 100 U/mL penicillin and 100 µg/mL streptomycin. Co-cultures were incubated at 37°C for up to 72 hours in an Incucyte S3 (Sartorius) with images collected every 4 hours. Growth of AML and ECs was monitored by detecting GFP or mKate signal, respectively. Cells were gated by size and GFP/mKate intensity and the aggregate intensity in each image was reported (GCU / RCU). The aggregate intensity of each well was normalized to the 0 hour time point (fold change) and plotted using Graphpad Prism software. For culture swap experiments, 10,000 target cells were co-cultured with CAR-T cells as described above. After 44 hours, plates were centrifuged at 300g for 5 minutes, supernatant was collected for cytokine analysis, and all remaining media was transferred to a second plate prepared with 10,000 THP-1 or TIME cells in 100 uL volume. Plates were centrifuged, all but ∼20 uL media was discarded and 180 uL of fresh media (RPMI for THP-1 and a 33% supplemented Vascular Cell Basal Media / 66% RPMI mix for TIME) was added to each well, then plates were returned to the Incucyte for an additional 48 hours.

#### ELISA

10,000 target cells (AML or endothelial cell lines) were co-cultured with CAR-T cells exactly as described above. After 24 hours, plates were centrifuged 5 minutes at 300g, then supernatant was collected, frozen at -20°C for long term storage, and analyzed for IFNγ and GzmB by ELISA (Biolegend) according to the manufacturer’s protocols.

### CAR Transduction Efficiency and CAR T Cell Phenotyping

100,000 target cells (THP-1 or TIME-mKate) were co-cultured with 50,000 CAR-T cells in a final volume of 1 mL RPMI media (for THP-1) or 1mL VCB media + 0.5 mL RPMI media (for TIME). After 48- or 72-hours cells were collected, washed in PBS with 2% FBS, incubated with Fc block (BD Pharmingen), then stained. At the 48 hour timepoint, T cells were stained for VE-cadherin-and CD93-CARs, followed by CD4/CD8/PD-1/TIM-3/LAG-3 antibodies. At the 72 hour timepoint, T cells were stained for CAR expression with Goat anti-igG antibody, followed by CD3/CD4/CD8/PD-1/TIM-3/LAG-3 antibodies. Following antibody staining, cells were analyzed with an Attune NxT flow cytometer (Thermo Fisher) and data were analyzed with FlowJo software.

### Preparation of LumeNEXT devices

LumeNEXT devices were fabricated in polydimethylsiloxane (PDMS) using standard soft lithography technique as previously described (22), with several differences as described below. Before use, LumeNEXT devices were treated with 2% poly(ethyleneimine) for 10 min followed by a 0·4% glutaraldehyde treatment for 30 min. Devices were washed four times with sterile deionized water. High-density rat-tail collagen type 1 (Corning) was diluted to a final concentration of 4 mg/mL and added into the main chamber of the device. Following collagen addition, devices were polymerized at room temperature for 10 minutes and then transferred to 37°C for 1 h to allow collagen to polymerize fully, at which point the PDMS rod was removed from the device to create molded lumens. HUVEC cells (Lonza, C2519A) were trypsinized, resuspended in ECM media (Sciencell, 1001) at 20,000 cells/μL and seeded into the lumens (2 µL per lumen). HUVEC-filled lumens were incubated at 37°C for 2 h to allow for cell attachment.

### LumeNEXT Assays

LumeNEXT vessels were cultured for 48h before use. VC.PD1/CD93.28ζ, VC.Pdel/CD93.28ζ, or mock CAR-T cells were stained with Vybrant™ DiI Cell-Labeling Solution (Invitrogen, V22885) (1:250), resuspended in a 1:1 ECM:DMEM mixture, and introduced into the lumen at the indicated concentrations. Cells were incubated for 16 hours. Following incubation, the lumens were washed with culture medium and stained with a mixture of Propidium iodide (1:500), Calcein AM (5 µg/mL), and Hoechst (1:1000). Samples were imaged using confocal microscopy (Nikon AXR). Viable cells were identified by green fluorescence (Calcein AM), dead cells by red fluorescence (Propidium iodide), and CAR T cells by deep-red fluorescence, which is displayed in blue in the merged images. Because they were no longer adherent, most dead cells were washed away as dyes were flushed through the lumen. Confocal images were acquired as z-stacks at multiple focal planes and analyzed using FIJI software. To quantify endothelial lumen coverage, each lumen was divided into top and bottom regions. Z-stack projections of the green channel (were generated separately for the top and bottom regions using FIJI. Endothelial cell coverage was calculated based on percentage of Calcein AM staining within the lumen ROI and the average of the two values was reported as the lumen coverage for each device.

### Statistical Analysis

Statistical analyses were carried out using GraphPad Prism software. Graphs represent averaged data including multiple biological replicates unless stated otherwise. *P* values were calculated with statistical tests described in each corresponding figure legend. *P* < 0.05 was considered statistically significant, and *P* values are denoted with asterisks as follows: *P* > 0.05 = not significant (ns), *P < 0.05 = *, P < 0.01 = **, P < 0.001 = ***, P < 0.0001 = \*\*\*\**.

## Results

### Expression of CDH5 Predicts VE-Cadherin Endothelial Specificity for iCAR Development

We interrogated bulk RNASeq data comparing gene expression in two endothelial cell (EC) lines (iHUVEC, TIME) and three AML cell lines (Kasumi-1, NOMO-1, THP-1), with the goal of identifying EC-specific surface markers that could serve as iCAR ligands (Fig. 1A). Considering the inflammatory milieu anticipated with CAR T cell therapy, the optimal iCAR ligand would retain surface expression on pro-inflammatory cytokine-exposed ECs while not emerging on AML cells to prevent opportunity for CAR T cell escape. We identified 12 differentially expressed genes with predicted surface-level expression that were stably upregulated on ECs at rest and across various exposures to IFNγ and TNFα (Fig. 1B). Among these genes, three encoded well-characterized EC proteins amenable to further study (*CDH5*, VE-cadherin; *KDR*, VEGFR2; *TEK*, Tie2). As predicted by the filtering strategy, transcript levels for these genes remained high in ECs but were not detected in AML cells (Fig. 1C). Cross-referencing expression of putative iCAR ligands with RNA expression patterns in the healthy single cell reference atlas Tabula Sapiens confirmed expression in ECs while redemonstrating the expression of *CD93* in ECs (Fig. 1D) (17). Like the expression of *CD93*, expression of these targets was observed across a diversity of organ-specific ECs in the healthy cell atlas, and present in vital organs enriched for *CD93* such as respiratory system and lung (Supp. Fig. 1A-B). By contrast, a large database of transcriptomic data from patient-derived AML samples (TARGET-AML, N=3,225) indicated that iCAR targets including *CDH5*, *KDR*, and *TEK* were not expressed or expressed at low levels in the vast majority of patients, whereas *CD93* was expressed at levels similar to or exceeding other AML immunotherapy markers in development (Fig. 1E, Supp. Fig. 1C). Based on this cross-referencing with large transcriptomic databases, we opted to pursue VE-cadherin and VEGFR2 as possible iCAR ligands.

**Figure 1:**
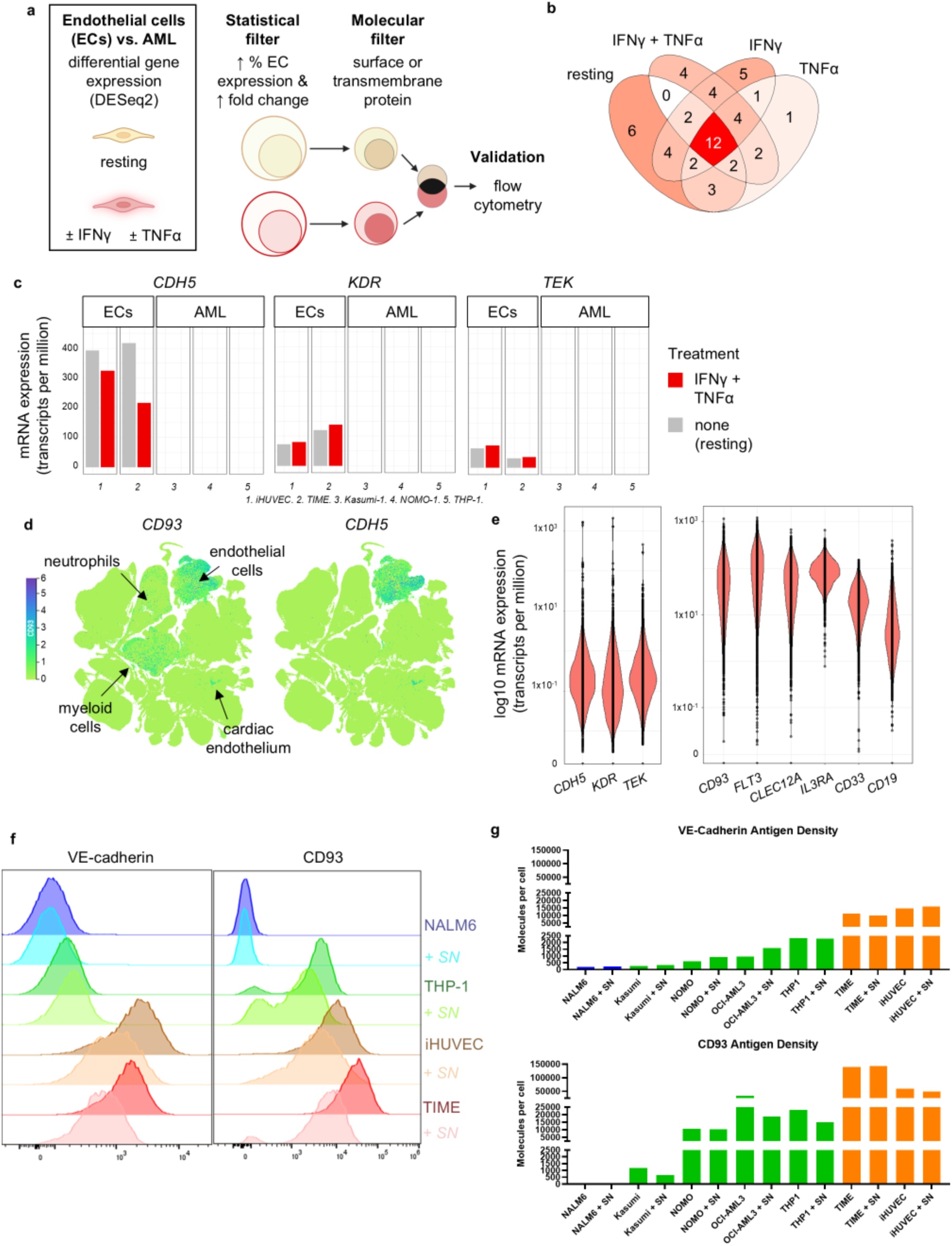
Bioinformatic approach to iCAR ligand selection. **A,** Schema for identifying candidate iCAR ligands among genes that are differentially expressed in endothelial cells (iHUVEC, TIME) as compared to acute myeloid leukemia (AML) cells (Kasumi-1, NOMO-1, THP-1), both at rest and with cytokine stimulation (IFNγ, TNFα, or both IFNγ and TNFα at 10 ng/mL for 24 hours). **B,** Venn diagram for genes upregulated on endothelial cells in resting and cytokine-activated states. Statistical threshold: FDR <0.05 and log2 fold change >10. Molecular threshold: annotation with “surface” or “transmembrane” protein in *in silico* human surfaceome (Wollschied lab). **C,** Expression levels (transcripts per million) of leading iCAR ligand candidates, at rest and with cytokine activation (IFNγ and TNFα; IFNγ only and TNFα only not shown). **D,** Expression of *CD93* is seen in both myeloid cells and endothelial cells within a large single cell atlas from healthy donors, Tabula Sapiens. *CDH5* expression is restricted to endothelial cells. **E,** Expression of iCAR ligands including *CDH5* is very low among 3,225 patient AML samples with transcriptomic data available on NCI GDC Data Portal. By contrast, *CD93* is almost universally expressed at high levels in patient AML samples, similar to or more than other AML immunotherapy markers. **F,** VE-cadherin (VC) and CD93 protein expression by flow cytometry on endothelial cell lines (iHUVEC, TIME) and on AML cell lines (THP-1), with B cell line (NALM6) as a negative control. Expression measured with and without 48 hours of exposure to supernatant (SN) from 24-hour co-culture of THP-1 cells and CD93 CAR T cells. **G,** VC and CD93 protein antigen density on endothelial cell lines (iHUVEC, TIME), AML cell lines (Kasumi-1, NOMO-1, OCI-AML3, THP-1), and a B cell line (NALM6) as a negative control. Expression measured with and without 24 hours of exposure to supernatant (SN) from 48-hour co-culture of THP-1 cells and CD93 CAR T cells at a 1:1 E:T ratio.

### VE-Cadherin Remains EC-Specific at the Protein Level Across Inflammatory Conditions

Using flow cytometry, we confirmed that VE-cadherin (VC) and VEGFR2 protein expression are specific to EC lines (iHUVEC, TIME) and lacking on AML cells (THP-1) (Fig. 1F). Consistent with our transcriptomic data, we observed that this EC specificity for VC persisted across inflammatory conditions, including after exposure to supernatant collected after co-culture of CD93 CAR T cells and AML cells. We also observed that VC antigen density on ECs remained high after exposure to supernatant from CAR T-AML interactions, a condition intended to mimic a pro-inflammatory microenvironment. In contrast, VEGFR2 expression decreased in the presence of this pro-inflammatory supernatant, limiting its potential usefulness as a CAR target (Fig. 1G, Supp. Fig. 1D) We therefore proceeded with VC as our iCAR ligand for the remainder of our experiments.

### A novel VE-Cadherin scFv Demonstrates Specificity for ECs

We engineered VC-directed specificity based on light and heavy chains from a murine anti-human VC antibody (2E11d), which had been developed previously in a study of EC adhesion junction function (20). Before inserting this VC-specific scFv into our NOT-gate system, we reasoned that incorporating it into a traditional CAR would establish specificity in the context of a CAR structure for cells expressing VC. Therefore, we first engineered a VC-specific aCAR, in either a light-heavy (LH) or heavy-light (HL) proximal-distal orientation, with a standard (G_4_S)_3_ linker connecting the two fragments, both linked to a CD28 hinge/transmembrane/costimulatory domain and CD3ζ activating domain (LH-VC.28ζ and HL-VC.28ζ, respectively) (Fig. 2A). Both VC.28ζ aCARs were transduced effectively, but HL-VC.28ζ was consistently expressed at higher levels across donors (Fig. 2B). Both LH-VC.28ζ and HL-VC.28ζ aCARs exhibited high specificity for endothelial cells as indicated by increases in cytokine production upon exposure of ECs but not AML cells (Fig. 2C, Supp. Fig. 2A). This EC specificity was also evident in the direct killing of ECs by both VC.28ζ.aCAR T cells (Fig. 2D, Supp. Fig 2B), whereas AML cell growth was equivalent between mock- and VC.28ζ CAR T-cell treated conditions (Fig. 2E, Supp. Fig 2C) Notably, IFNγ production and cytotoxicity, especially at lower E:T ratios, was significantly increased for VC.28ζ CARs as compared to CD93.28ζ CARs, which was somewhat unexpected given the high antigen density of both targets on the endothelial cell surface (Fig. 1G). We opted to use the HL rather than LH orientation for all future constructs given the higher HL CAR expression and similar capacity to discriminate between ECs and AML.

**Figure 2:**
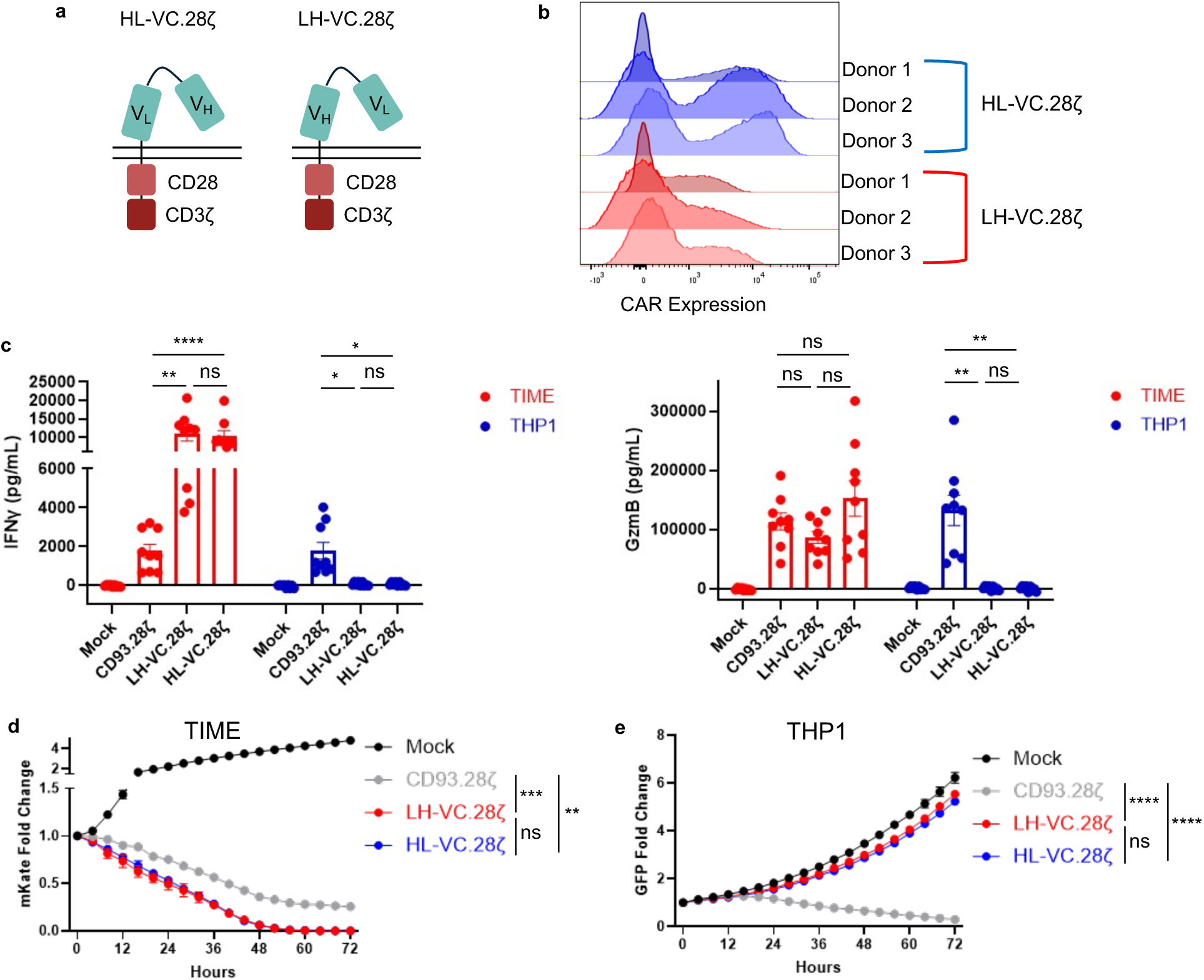
Validation of CARs containing a novel VE-Cadherin ScFv. **A,** Schema for VC aCARs, with either the scFv heavy chain distal to the light chain (HL) or vice versa (LH). V_H_, heavy chain variable fragment; V_L_, light chain variable fragment. **B,** Flow cytometry histograms confirming detection of VC scFv on VC aCAR T cells measured with anti-IgG antibody. **C,** Effector cytokine levels in supernatant from 24h co-culture of CAR T cells with ECs (TIME) or AML cells (THP-1) at 1:2 E:T ratio, n=3 (one-way ANOVA). Comparing CD93.28ζ and HL-VC.28ζ IFNγ production versus cell lines at 1:2 E:T ratio, p <0.0001 for TIME and p = 0.0025 for THP-1. Comparing CD93.28ζ and HL-VC.28ζ GzmB production versus cell lines at 1:2 E:T ratio, p = 0.8018 for TIME and p = 0.0044 for THP-1 **D,** Survival of ECs expressing mKate (TIME) over a 68h co-culture with CAR T cells at 1:2 E:T ratio, as measured by fluorescence in Incucyte killing assay. One representative donor is shown, with statistics based on summary data including n=3 separate donors (one-way ANOVA of AUC). **E,** Survival of AML expressing GFP (THP-1) over a 68h co-culture with CAR T cells at 1:2 E:T ratio, as measured by fluorescence in Incucyte killing assay. One representative donor is shown, with statistics based on summary data including n=3 separate donors (one-way ANOVA of AUC).

### Introduction of an iCAR has Minimal Impact on Baseline CD93 CAR T Cell Product Characteristics

We chose to design our VC-specific iCAR with the well-characterized inhibitory domain of PD-1, which has been previously incorporated into iCAR design for NOT-gated CAR T cells (9, 10). The VC-specific scFv was directly linked to the PD-1 intracellular domain via a CD8α hinge and transmembrane domain (VC.PD1), and a version lacking the intracellular signaling domain (VC.Pdel) was used as a control (Fig. 3A). The VC.PD1 and VC.Pdel iCARs were then cloned into a bicistronic vector upstream of the CD93.28ζ aCAR for retroviral transduction into healthy donor T cells (Fig. 3B). The resulting CAR T cells co-expressed aCAR and iCAR constructs at a near 1:1 ratio (Fig. 3C). The mean fluorescence intensities of both the CD93 aCAR and the VC-specific iCAR were slightly lower in the Pdel control compared to the full-length PD-1 iCAR (Fig. 3D-E). We did not observe any significant differences in CAR T cell proliferation during the 10 day production and expansion period, even when compared to CD93.28z CAR T cells without any iCAR (Supp. Fig 3A). Similarly, introducing the PD1 iCAR construct did not significantly affect baseline expression of native PD-1 or other markers associated with T-cell exhaustion, such as TIM3 and LAG3, as compared to the Pdel control in either the total CAR+ or CD4+ / CD8+ CAR+ populations (Fig. 3F-G, Supp. Fig 3B-D) (23).

**Figure 3:**
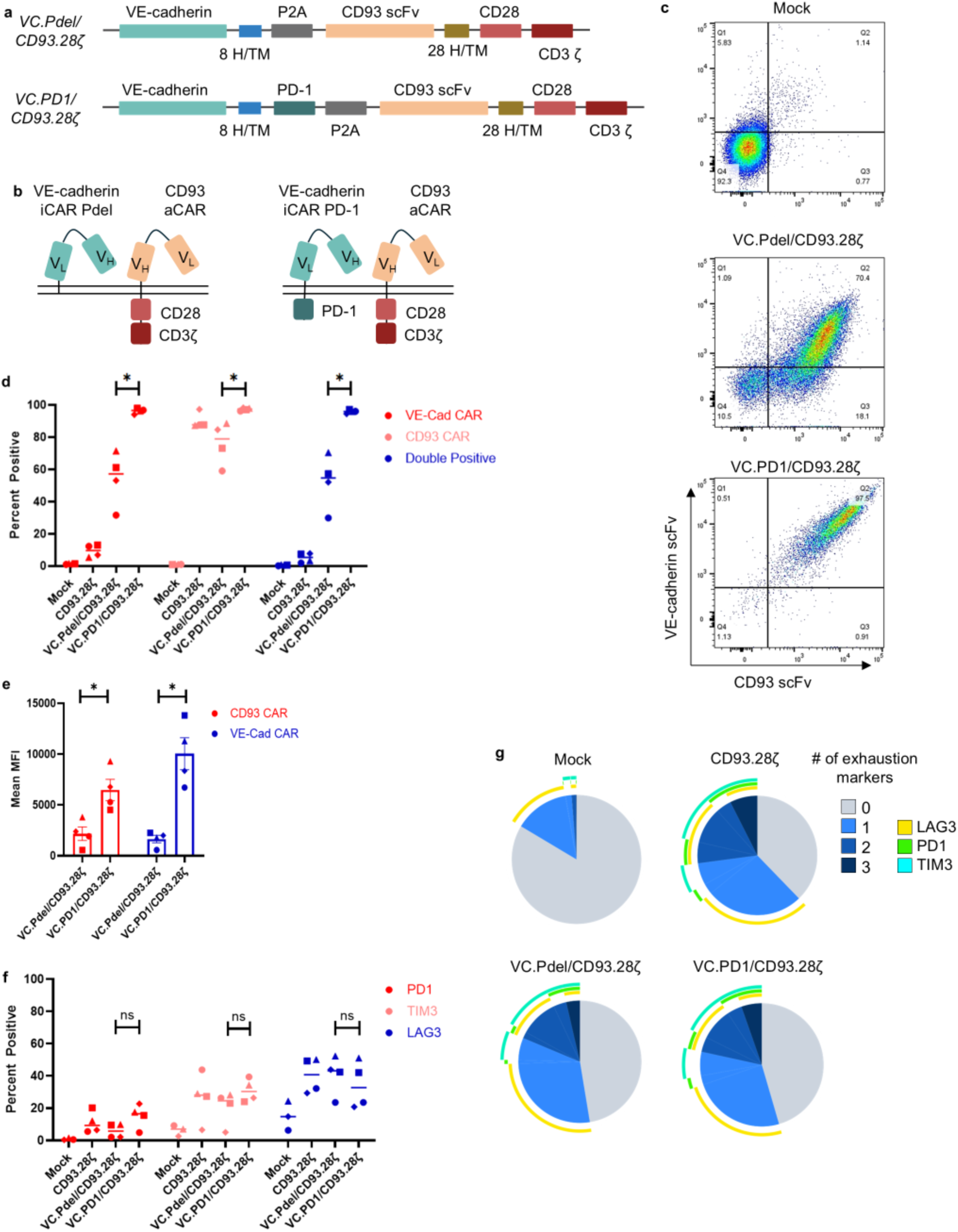
Characterization of NOT-gated CAR T cells. **A,** Plasmid for bicistronic vector including VC scFv adjoined to PD-1 inhibitory domain (or Pdel control). **B**, Schema for bicistronic vector gene products: 1) VC Pdel (control) iCAR with CD93 aCAR (VC.Pdel/CD93.28ζ); 2) VC PD-1 iCAR with CD93 aCAR (VC.PD1/CD93.28ζ). **C,** Flow cytometry confirming co-expression of retroviral vector gene products on T cells from one representative donor, as measured by detection of VC or CD93 scFv. **D,** Percent positivity of CD93 aCAR or VC iCAR across four separate donors (unpaired t-test). Comparing VC.Pdel with VC.PD1 VC CAR positivity, p=0.0142. Comparing VC.Pdel with VC.PD1 CD93 CAR positivity, p= 0.0475. Comparing VC.Pdel with VC.PD1 double positivity, p = 0.0131. Right, MFI of CD93 aCAR or VC aCAR among double positive CAR T cells. Comparing CD93 CAR MFI p = 0.0178. Comparing VC CAR MFI p = 0.0106 (unpaired t-test). **E,** Median Fluorescence Intensity (MFI) of VE-Cadherin and CD93 CAR within the double positive populations as shown in Figure 2C (unpaired t-test). **F**, Baseline expression of PD-1, TIM3, and LAG3 within CAR+ populations, as gated on FMO controls (unpaired t-test). **G,** SPICE plots of T cell exhaustion markers (PD1, TIM-3 and LAG-3) measured at baseline by flow cytometry (29). One representative donor is shown.

### VC/CD93 NOT-Gated CAR T Cells Kill AML Cells and Spare ECs

We proceeded to test whether cytokine production and cytotoxicity differed with exposure to AML cells and ECs for these CAR T cell variants. VC.Pdel/CD93.28ζ CAR T cells produced cytokines associated with T cell effector function (IFNγ and GzmB) when exposed to AML (THP-1 and OCI-AML3) as well as ECs (TIME and iHUVEC), as expected given the fully functional CD93 aCAR and VC iCAR lacking a functional PD-1 inhibitory domain (Fig. 4A, Supp. Fig. 4A). By contrast, VC.PD1/CD93.28ζ CAR T cells secreted cytokines in response to AML, but this activity was almost completely abrogated when these cells were co-cultured with ECs instead, indicating EC-specific CAR T-cell inhibition dependent on PD-1. Similarly, CD93.28ζ/VC.PD1 CAR T cells killed AML cells but not ECs in live cell cytotoxicity assays (Fig. 4B, Supp. Fig. 4B-C). These results showed that VC/CD93 NOT-gated CAR T cells can provide tumor specificity and spare ECs in an *in vitro,* two-dimensional co-culture system.

**Figure 4:**
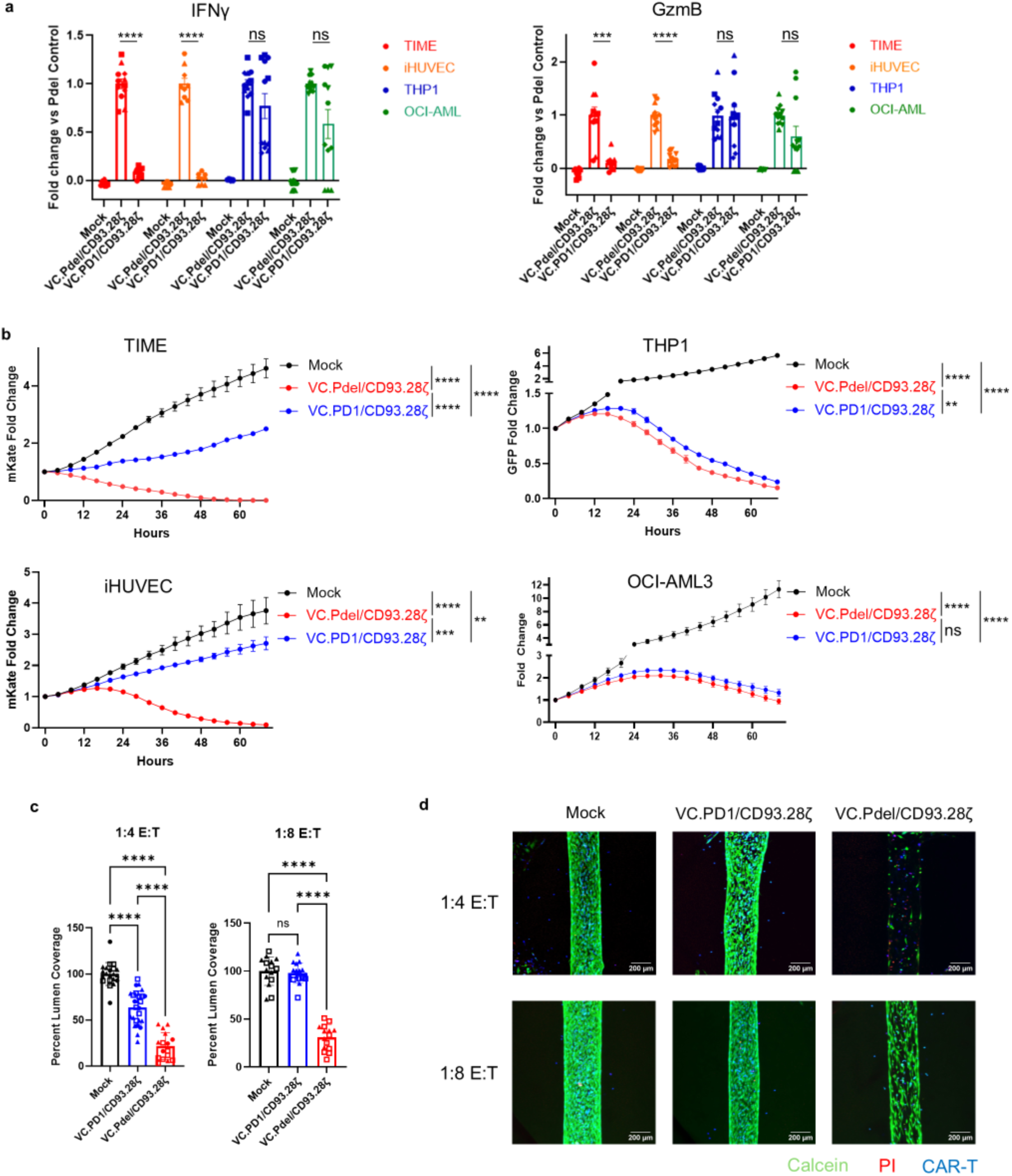
Functional validation and AML specificity of NOT-gated CAR T cells. **A,** Effector cytokine levels in supernatant from 24h co-culture of CAR T cells with ECs (TIME, iHUVEC) or AML cells (THP-1, OCI) at a 1:2 E:T ratio. Cytokine levels depicted as fold change compared to Pdel control, normalized on each assay date (one-way ANOVA). Individual donors are represented with different shapes (n=4 donors). Comparing VC.PD1/CD93.28ζand VC.Pdel/CD93.28ζ IFNγ versus TIME and iHUVEC, p <0.0001. Comparing VC.PD1/CD93.28ζand VC.Pdel/CD93.28ζ GzmB versus TIME, p = 0.0005. Comparing VC.PD1/CD93.28ζand VC.Pdel/CD93.28ζ GzmB versus iHUVEC, p <0.0001. **B,** Survival of either ECs expressing mKate (TIME, iHUVEC) or AML expressing GFP (THP-1, OCI-AML3) over a 68h co-culture with CAR T cells at 1:2 E:T ratio, as measured by fluorescence in Incucyte killing assay. One representative donor is shown for each target cell line, with significance bars showing summary data including n=3 separate donors (one-way ANOVA of AUC). Comparing VC.PD1/CD93.28ζand VC.Pdel/CD93.28ζ versus TIME, p <0.0001. Comparing VC.PD1/CD93.28ζand VC.Pdel/CD93.28ζ versus iHUVEC, p = 0.0001. Comparing VC.PD1/CD93.28ζand VC.Pdel/CD93.28ζ GzmB versus THP-1, p = 0.0027. **C,** LumeNEXT assays confirm NOT-gate inhibition in a three-dimensional model. Lumens seeded with HUVEC cells were co-cultured with labeled T cells for 16-hours at varying ratios, then labeled with Propidium Iodide (PI) and calcein dye, then imaged by confocal microscopy. Percent lumen coverage was measured (one-way ANOVA). Individual donors are represented with different shapes (n= 2-3 donors). Comparing VC.PD1/CD93.28ζ and VC.Pdel/CD93.28ζ at 1:4 E:T and 1:8 E:T ratios, p <0.0001 **D,** Representative images of the LumeNEXT assay from a single donor. Live HUVEC cells are labeled in green, dead cells in red, and T cells in blue.

We then tested the inhibition capacity of the NOT-Gated CAR T cells in the LumeNEXT platform, a microfluidic-based method that recapitulates a three-dimensional endothelialized luminal structure and mimics the native geometry of blood vessels, including luminal-basal polarity (22). Across a range of effector-to-target (E:T) ratios, VC.PD1/CD93.28ζ CAR T cells consistently displayed reduced cytotoxicity relative to VC.Pdel/CD93.28ζ controls, providing strong functional validation that NOT-gating preserves endothelial cell integrity within native vascular structures (Fig. 4C-D, Supp. Fig 4D-E).

To determine whether the discriminatory capacity of NOT-gated CAR T cells was related to target cell-induced CAR T cell exhaustion or dysfunction, we compared expression of native PD-1, LAG3, and TIM3 on VC.PD1/CD93.28ζ and VC.Pdel/CD93.28ζ CAR T cells after a 3-day co-culture with AML or endothelial cells. As expected, expression of all three markers was more prevalent on CAR T cells after co-culture (Supp. Fig 5A-B) compared to baseline (Fig. 3G, Supp. Fig 3B), but there were no significant differences among CD93 CAR T cells, for the whole population or among CD4+ or CD8+ sub-populations (Supp. Fig 5C-E). No significant changes in CD4-8 ratio were observed between VC.PD1/CD93.28ζ and VC.Pdel/CD93.28ζ CAR T cells following either AML or EC co-culture (Supp. Fig 5F).

**Figure 5:**
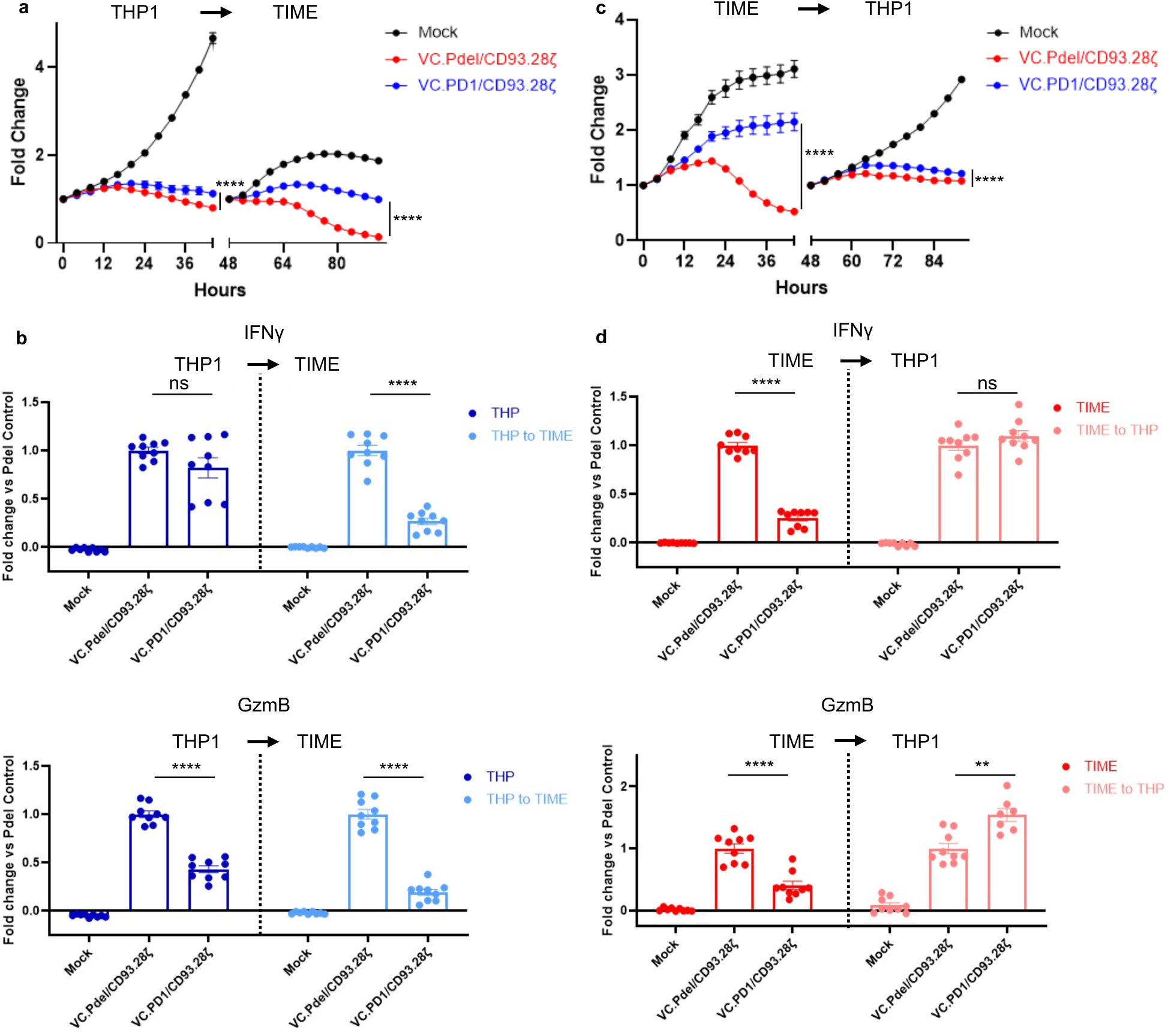
NOT-gated functions are reversible. **A,** Survival of either ECs expressing mKate (TIME, iHUVEC) or AML expressing GFP (THP-1, OCI-AML3), as measured by fluorescence in Incucyte killing assay CAR T cells were first co-cultured with THP-1 for 44 hours at 1:2 E:T ratio, then transferred to a plate seeded with TIME cells for an additional 44 hours. One representative donor is shown, with statistics based on summary data including n=3 separate donors (one-way ANOVA of AUC). Comparing VC.PD1/CD93.28ζ and VC.Pdel/CD93.28ζ at 44- and 92-hour timepoints, p <0.0001. **B,** Effector cytokine levels in supernatants taken at 44- and 92-hour timepoints of co-culture with THP-1 or TIME cells at a 1:2 E:T ratio. Cytokine levels depicted as fold change compared to Pdel control, normalized on each assay date (one-way ANOVA). Comparing VC.PD1/CD93.28ζ and VC.Pdel/CD93.28ζ IFNγ and GzmB at the 92-hour timepoint, p <0.0001. Comparing VC.PD1/CD93.28ζ and VC.Pdel/CD93.28ζ GzmB at the 44 hour timepoint, p <0.0001.**C,** Survival of either ECs expressing mKate (TIME, iHUVEC) or AML expressing GFP (THP-1, OCI-AML3), as measured by fluorescence in an Incucyte killing assay. CAR T cells were first co-cultured with TIME for 44 hours at 1:2 E:T ratio, then transferred to a plate seeded with THP-1 for an additional 44 hours. One representative donor is shown, with statistics based on summary data including n=3 separate donors (one-way ANOVA of AUC). Comparing VC.PD1/CD93.28ζ and VC.Pdel/CD93.28ζ at 44- and 92-hour timepoints, p <0.0001. **D,** Effector cytokine levels in supernatants taken at 44- and 92-hour timepoints of co-culture with THP-1 or TIME cells at a 1:2 E:T ratio. Cytokine levels depicted as fold change compared to Pdel control, normalized on each assay date (one-way ANOVA). Comparing VC.PD1/CD93.28ζ and VC.Pdel/CD93.28ζ IFNγ and GzmB at the 44-hour timepoint, p <0.0001. Comparing VC.PD1/CD93.28ζ and VC.Pdel/CD93.28ζ GzmB at the 92-hour timepoint p = 0.0044.

### Inhibition and Activation of NOT-Gated CAR T Cells are Reversible

We next evaluated whether prior CAR T cell activation with AML killing might influence the VC iCAR-mediated protection of ECs, and conversely whether prior CAR T cell inhibition would prevent desired AML recognition and cytotoxicity, thus modeling the more complex cellular encounters over time that these CAR T cells would have in a physiologic system. To assess the dynamics of target-specific activation and inhibitory signals, we exposed NOT-gated CAR T cells in series either to AML then ECs, or to ECs then AML. Critically, initial exposure to AML did not abrogate the ability of VC.PD1/CD93.28ζ CAR T to protect ECs (Fig. 5A-B, Supp. Fig. 6A-B), whereas VC.Pdel/CD93.28ζ CAR T cells still exhibited toxicity to ECs after transfer. Conversely, VC.PD1/CD93.28ζCAR T cells that first protected ECs still effectively killed AML cells upon transfer (Fig. 5C-D, Supp. Fig. 6C-D). Compared to mock, NOT-gated CAR T cells did limit endothelial cell growth to a greater extent with the donors used in this experiment compared to Fig. 3, but they were still largely protected from cytotoxicity with stable growth curves regardless of the order of target cell exposure. Together, these results indicate that the NOT-gated CAR T cells can turn “on” and “off” as predicted by their target cell engagement, regardless of prior exposures.

## Discussion

Our study demonstrates that VC/CD93 NOT-gated CAR T cells effectively kill AML *in vitro* while sparing CD93+ ECs at risk for OTOT toxicity. Importantly, the EC protection afforded by the VC iCAR is not abrogated by prior activation of CAR T cells encountering AML cells. This sustained NOT-gate function is the foundation for further developing a safe and effective CD93-directed AML immunotherapy. Our bioinformatics-guided target identification strategy is also broadly applicable as an approach to OTOT toxicity problems that may limit cell therapy options for other difficult-to-treat malignancies. Selecting iCAR candidates that are predicted to remain specific to healthy tissue despite a changing inflammatory milieu can help narrow candidate iCAR targets and was a successful strategy here. It adds to the existing principles proposed by others to guide combinatorial CAR pairing, both for the selection of iCAR ligands and for choosing other tumor aCAR targets with alternative logic-gating strategies, such as AND gating (24). For AML in particular, there are also data that pro-inflammatory cytokines have a direct influence on leukemic survival during CAR T-cell therapy independent of target antigen expression, further supporting pre-clinical work that incorporates the cytokine environment as an essential variable (25). We were struck by the superior protection afforded to ECs using the VC iCAR ligand as compared to CD19 as a surrogate EC-specific marker in our previous work (9). It seems possible that the native proximity of VC and CD93 within EC junctions, where CD93 has been shown to stabilize VC turnover (26), may increase the likelihood that a NOT-gated CAR T cell receives VC-specific inhibitory signaling anytime it binds nearby CD93. Accordingly, spatial proximity of aCAR and iCAR ligands may be a relevant design principle for further applications of NOT-gating.

Our NOT-gated CAR T cells demonstrated a distinctive ability to transition between activated and inhibitory states based on target cell engagement. While an essential safety feature, this agility is not a foregone conclusion with increasingly complex T-cell receptor signaling. Within non-transduced T cells, for example, PD-1 expression inhibits T cell proliferation and cytokine signaling to a degree that exceeds what is predicted by ligand-binding affinity, and PD-1 can suppress subsequent antigen recognition by native T-cell receptors (TCR) (27). The reverse does not automatically hold true, and indeed strong TCR activation may not override subsequent inhibitory signals but rather lead to direct T cell inhibition via intrinsic negative feedback loops (28). Our CAR T cells did not seem to be affected by the same inhibitory bias, perhaps because CARs do not access the full regulatory machinery of a native TCR.

In summary, our VC/CD93 NOT-gated CAR T cells successfully distinguish ECs from AML *in vitro*. A bioinformatics-guided approach can efficiently identify iCAR ligands, which may be further selected based on stable expression during inflammation and predicted spatial proximity to the aCAR ligand. This strategy may advance CAR T cell therapies for AML or other malignancies where OTOT toxicity is limiting.

## DECLARATIONS

### Ethics approval and consent to participate

Not applicable

### Consent for publication

Not applicable

### Availability of data and material

All data relevant to the study are included in the article or uploaded as supplementary information. Bulk RNAseq data from this study are available in the NCBI GEO database (GSE159991).

### Authors contributions

TW and OAK collected data and contributed to assay design, data interpretation, and manuscript preparation. KS, CS, PV, and LM collected data and contributed to assay design and data interpretation. SCK and RMR oversaw experimental design and data interpretation, and finalized the manuscript. All authors approved the final version of the manuscript.

## Acknowledgements

We would like to acknowledge Dr. Barry Gumbiner (University of Virginia) for kindly providing the sequence from which the VE-cadherin scFv was generated. We are grateful to Dr. Crystal Mackall, Ravi Majeti, and Elena Sotillo (Stanford) for providing cell lines and for their mentorship. We would like to thank Lance Rodenkirch and the UW Optical Imaging Core (UWOIC) for their help performing LumeNEXT experiments. We would also like to thank Dr David Beebe for his mentorship. Some of these data were presented at the 2025 Society of Immunotherapy for Cancer Annual Meeting. The graphical abstract and illustrations in Figures 1,2, and 3 were created with Biorender.com.

## LIST OF ABBREVIATIONS

AML: Acute myeloid leukemia
aCAR: Activating chimeric antigen receptor
B-ALL: B-cell acute lymphoblastic leukemia
CAR: Chimeric antigen receptor
EC: Endothelial cell
ELISA: Enzyme-linked immunosorbent assay
E:T ratio: Effector-to-target ratio
GFP: Green fluorescent protein
iCAR: Inhibitory chimeric antigen receptor
IFNγ: Interferon-gamma
iHUVEC: Immortalized human umbilical vein endothelial cells
HSCT: Hematopoietic stem cell transplant
HSPC: Hematopoietic stem and progenitor cell
LSC: Leukemic stem cell
MHC: Major histocompatibility complex
OTOT: On-target, off-tumor
scFv: Single chain variable fragment
TIME: Telomerase-immortalized microvascular endothelial cells
TNFα: Tumor necrosis factor-alpha
VC: VE-cadherin
VEGFR2: Vascular endothelial growth factor receptor 2

