## Supplemental Figures for "VE-cadherin NOT-gated CD93 CAR T cells discriminate between AML and healthy endothelial cells"

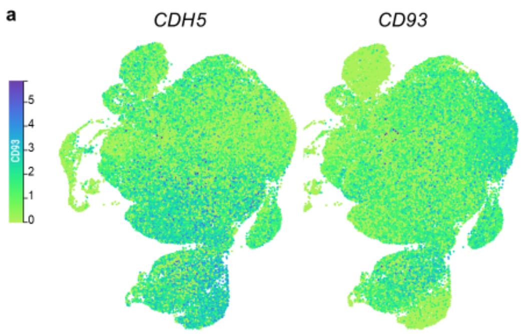


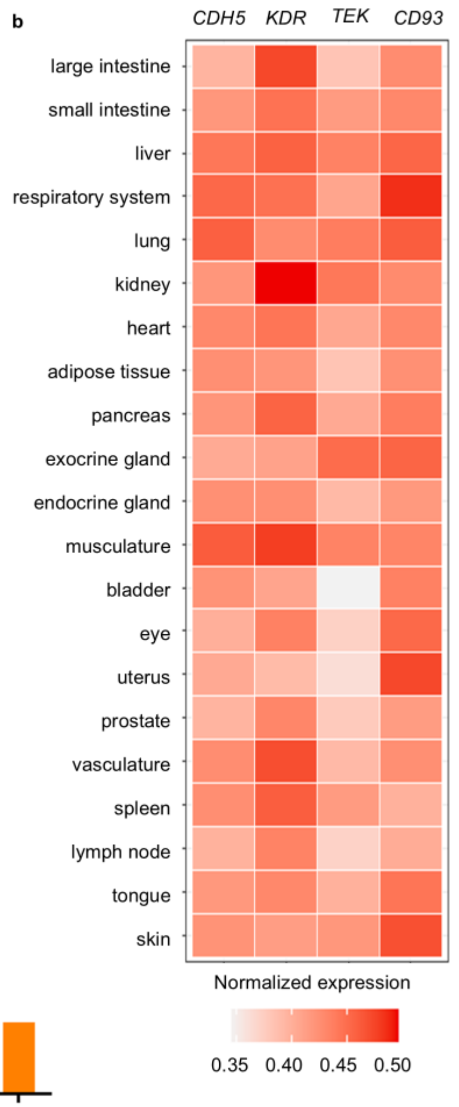


c

| Gene | Protein | Expression *median (IQR)* |
|---|---|---|
| *CDH5* | VE-cadherin | 0.17 (0.07-0.43) |
| *KDR* | VEGFR2 | 0.08 (0.02-0.32) |
| *TEK* | Tie2 | 0.19 (0.07-0.51) |
| *CD93* | CD93 | 36.7 (11.5-89.2) |
| *FLT3* | FLT3 | 65.2 (20.0-163) |
| *CLEC12A* | CLL-1 | 37.8 (13.7-85.1) |
| *IL3RA* | CD123 | 81.4 (53.8-121) |
| *CD33* | CD33 | 17.2 (8.5-31.2) |
| *CD19* | CD19 | 4.2 (1.83-10.8) |

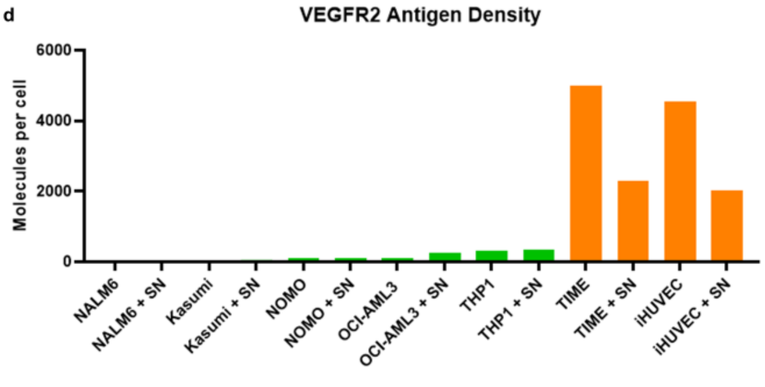


**Supplement 1. A,** Expression of both *CD93* and *CDH5* spans multiple organ-specific endothelial cell populations in Tabula Sapiens. **B,** Normalized transcript expression of iCAR ligands (*CDH5*, *KDR*, *TEK*) and *CD93* in endothelial cell subsets in Tabula Sapiens, coded by organ of origin. **C ,** Median transcript expression for possible iCAR ligands and AML immunotherapy targets among 3,225 patient AML samples with transcriptomic data available on NCI GDC Data Portal. IQR, interquartile range. **D,** VEGFR2 protein antigen density on endothelial cell lines (iHUVEC, TIME), AML cell lines (Kasumi-1, NOMO-1, OCI-AML3, THP-1), and a B cell line (NALM6) as a negative control. Expression measured with and without 24 hours of exposure to supernatant (SN) from 48-hour co-culture of THP-1 cells and CD93 CAR T cells at a 1:1 E:T ratio.

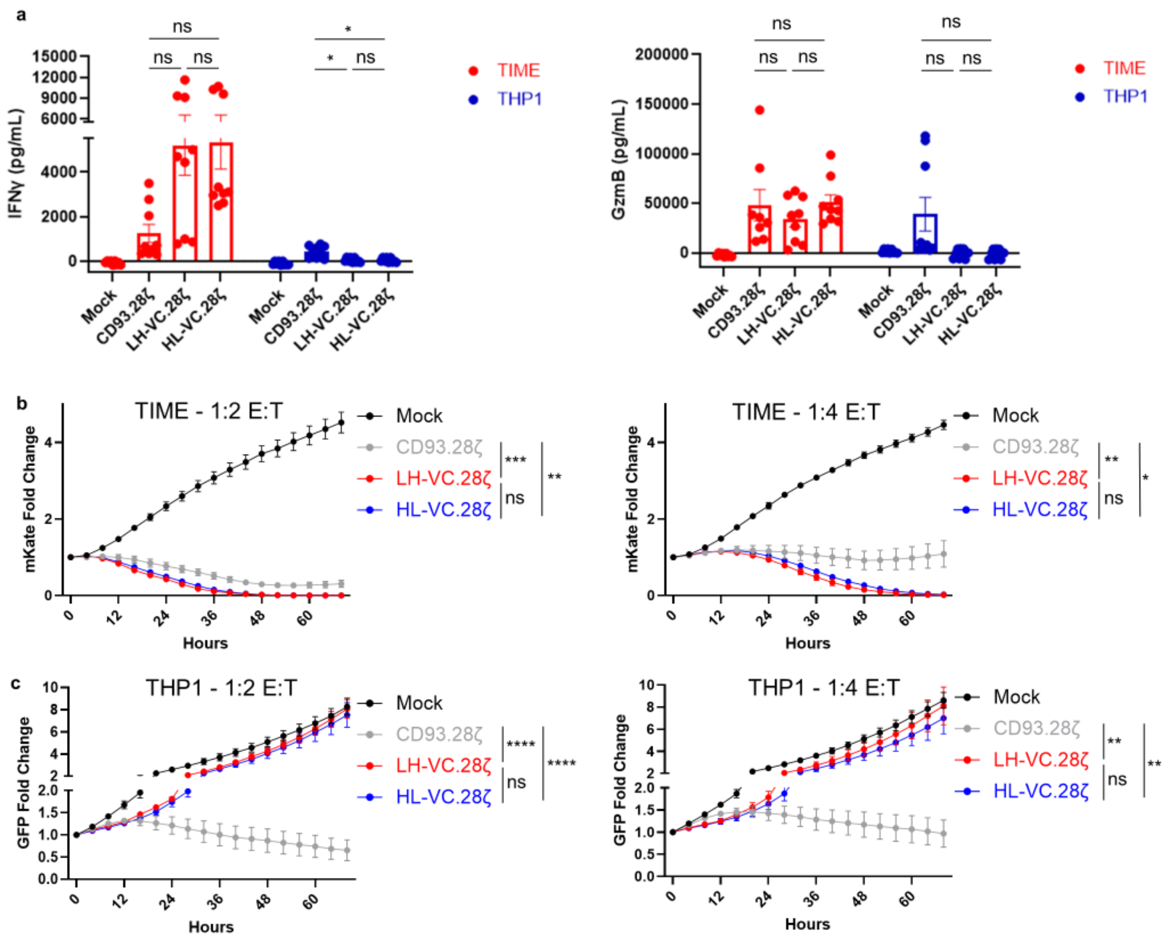


**Supplement 2. A,** Effector cytokine levels in supernatant from 24h co-culture of CAR T cells with ECs (TIME) or AML cells (THP-1) at 1:4 E:T ratio, n=3 donors (one-way ANOVA). **B,** Summary Incucyte data of CAR T and TIME co-culture from Fig 3B at 1:2 and 1:4 E:T ratios. Includes n=3 donors (one-way ANOVA of AUC). **C,** Summary Incucyte data of CAR-T and THP-1 co-culture from Fig 3B at 1:2 and 1:4 E:T ratios. Includes n=3 donors (one-way ANOVA of AUC).

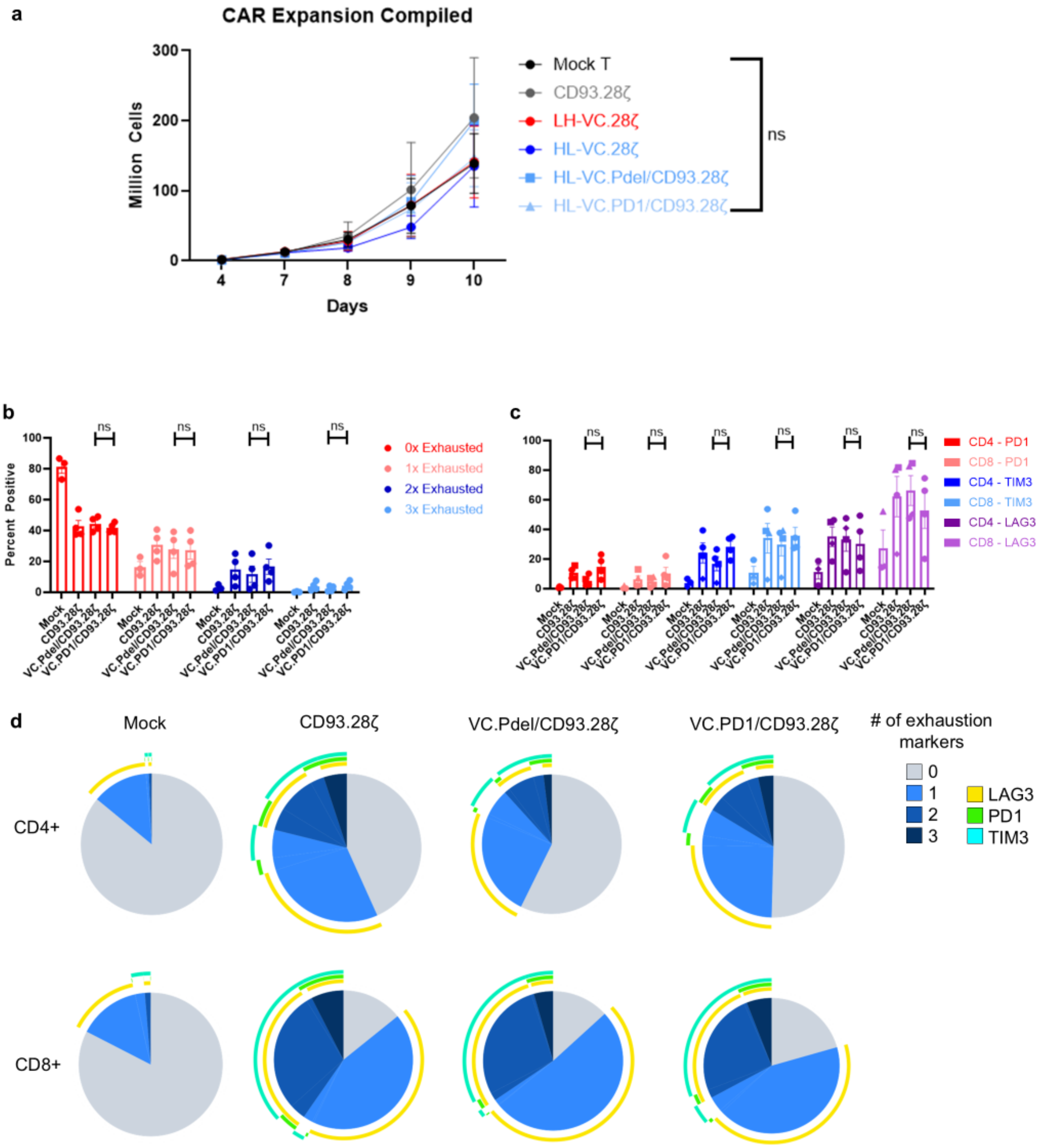


**Supplement 3. A,** Expansion of T cells was monitored over time. CAR-T cells were transduced on days 2-3 after activation, then CD3/28 beads were removed on day 4. Summary data of n= 3-4 replicates shows no significant difference between any T cell populations (one-way ANOVA). **B,** Cumulative expression of T cell exhaustion markers (PD1, TIM-3 and LAG-3) measured at baseline by flow cytometry. Summary data of n=3 donors is shown. No significant difference in exhaustion of VC.PD1/CD93.28ζ and VC.Pdel/CD93.28ζ was observed (unpaired T-test). **C,** Baseline expression of PD-1, TIM3, and LAG3 within CAR+ populations, separated into CD4+ and CD8+ populations (unpaired t-test). **D,** SPICE plots of T cell exhaustion markers (PD1, TIM-3 and LAG-3) measured at baseline by flow cytometry {Roederer, 2011 #275}. One representative donor is shown. CD4+ and CD8+ subpopulations were separately plotted.

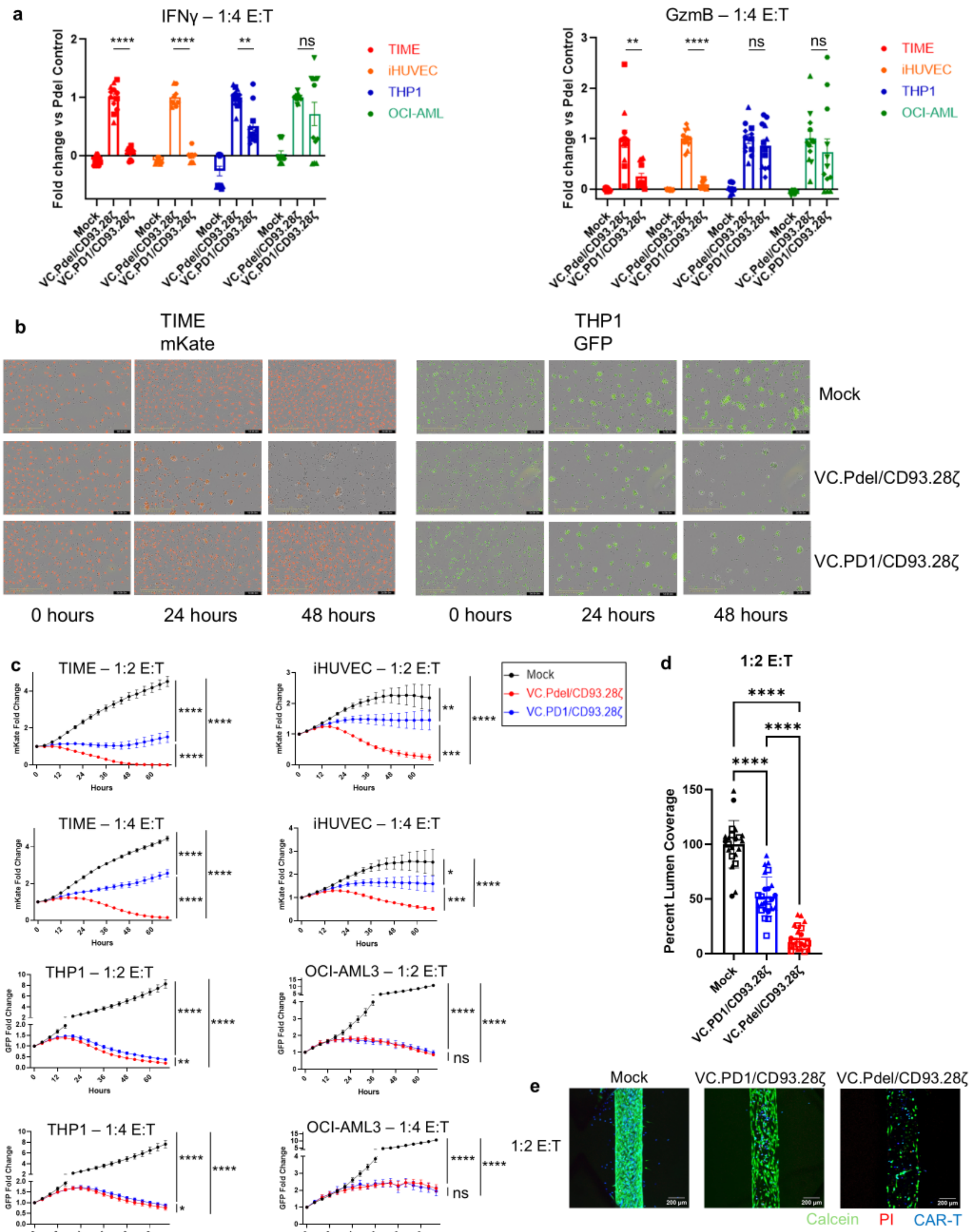


**Supplement 4. A,** Effector cytokine levels in supernatant from 24h co-culture of CAR T cells with ECs (TIME, iHUVEC) or AML cells (THP-1, OCI) at a 1:4 E:T ratio. Cytokine levels depicted as fold change compared to Pdel control, normalized on each assay date (one-way ANOVA). Individual donors are represented with different shapes (n=4 donors). **B,** Representative Incucyte images of CAR T cell co-cultures with either TIME mKate or THP1 GFP cells. Three

timepoints are shown of each well (0/24/48 hours) containing either mock T, VC.Pdel/CD93.28ζ or VC.PD1/CD93.28ζ at a 1:2 E:T ratio. **C,** Summary Incucyte data from Figure 3B. **D,** Summary LumeNEXT data from n= 3 donors at a 1:2 E:T ratio is shown (One-way ANOVA of AUC). Percent lumen coverage was measured (one-way ANOVA). Individual donors are represented with different shapes (n=3 donors). Comparing VC.PD1/CD93.28ζ and VC.Pdel/CD93.28ζ at 1:2 E:T ratio, $p < 0.0001$. **E,** Representative images of the LumeNEXT assay from a single donor. Healthy HUVEC cells are labeled in green, dying cells in red, and T cells in blue.

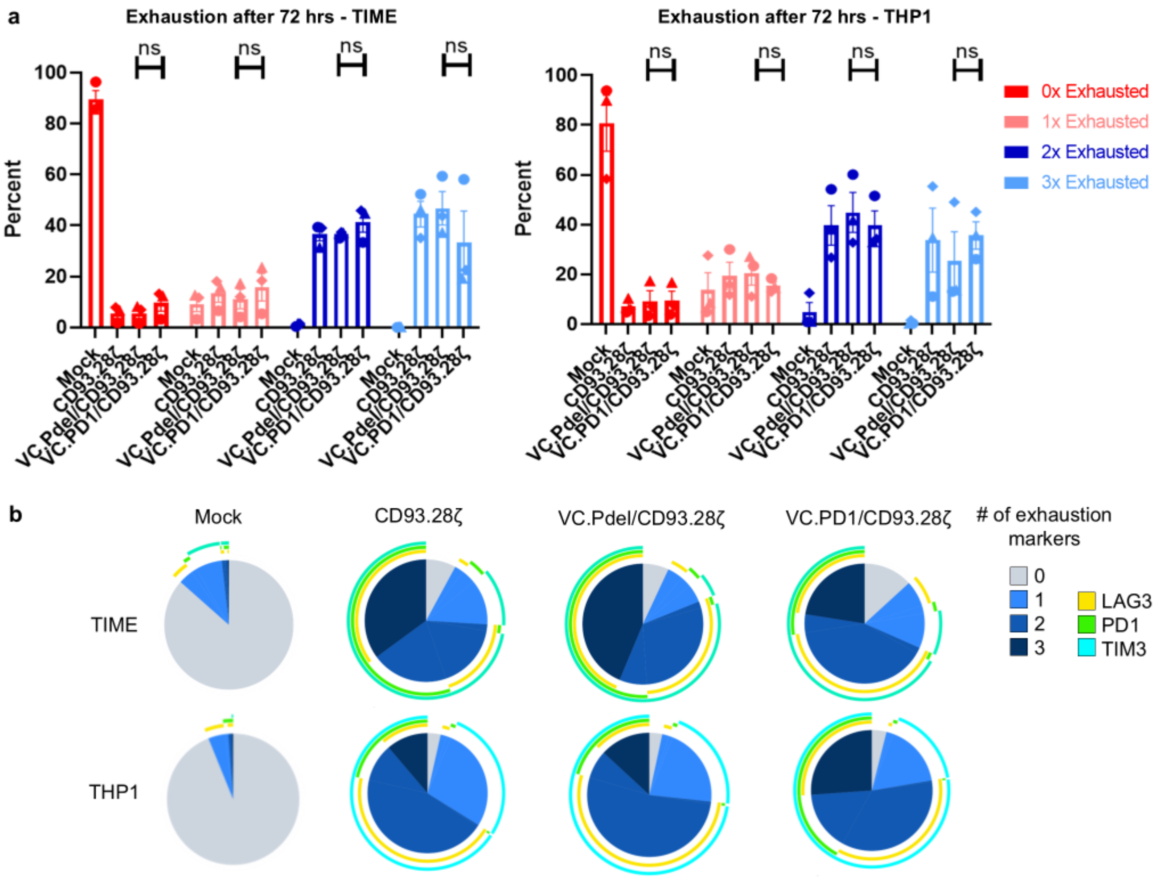
a
Exhaustion after 72 hrs - TIME
Exhaustion after 72 hrs - THP1
ns
Percent
0
20
40
60
80
100
Mock
CD93.28ζ
VC.Pdel/CD93.28ζ
VC.PD1/CD93.28ζ
0x Exhausted
1x Exhausted
2x Exhausted
3x Exhausted
b
Mock
CD93.28ζ
VC.Pdel/CD93.28ζ
VC.PD1/CD93.28ζ
TIME
THP1
### of exhaustion markers
0
1
2
3
LAG3
PD1
TIM3

c

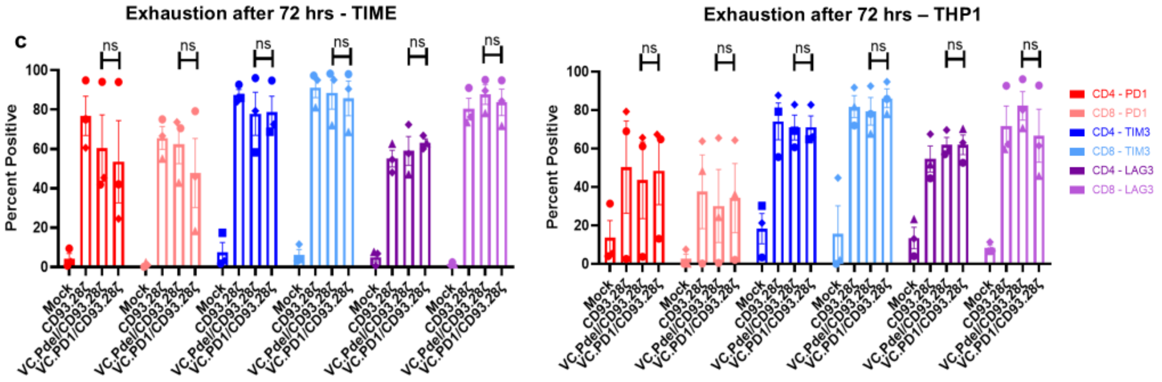
Exhaustion after 72 hrs - TIME
Exhaustion after 72 hrs – THP1
Percent Positive
CD4 - PD1
CD8 - PD1
CD4 - TIM3
CD8 - TIM3
CD4 - LAG3
CD8 - LAG3

d

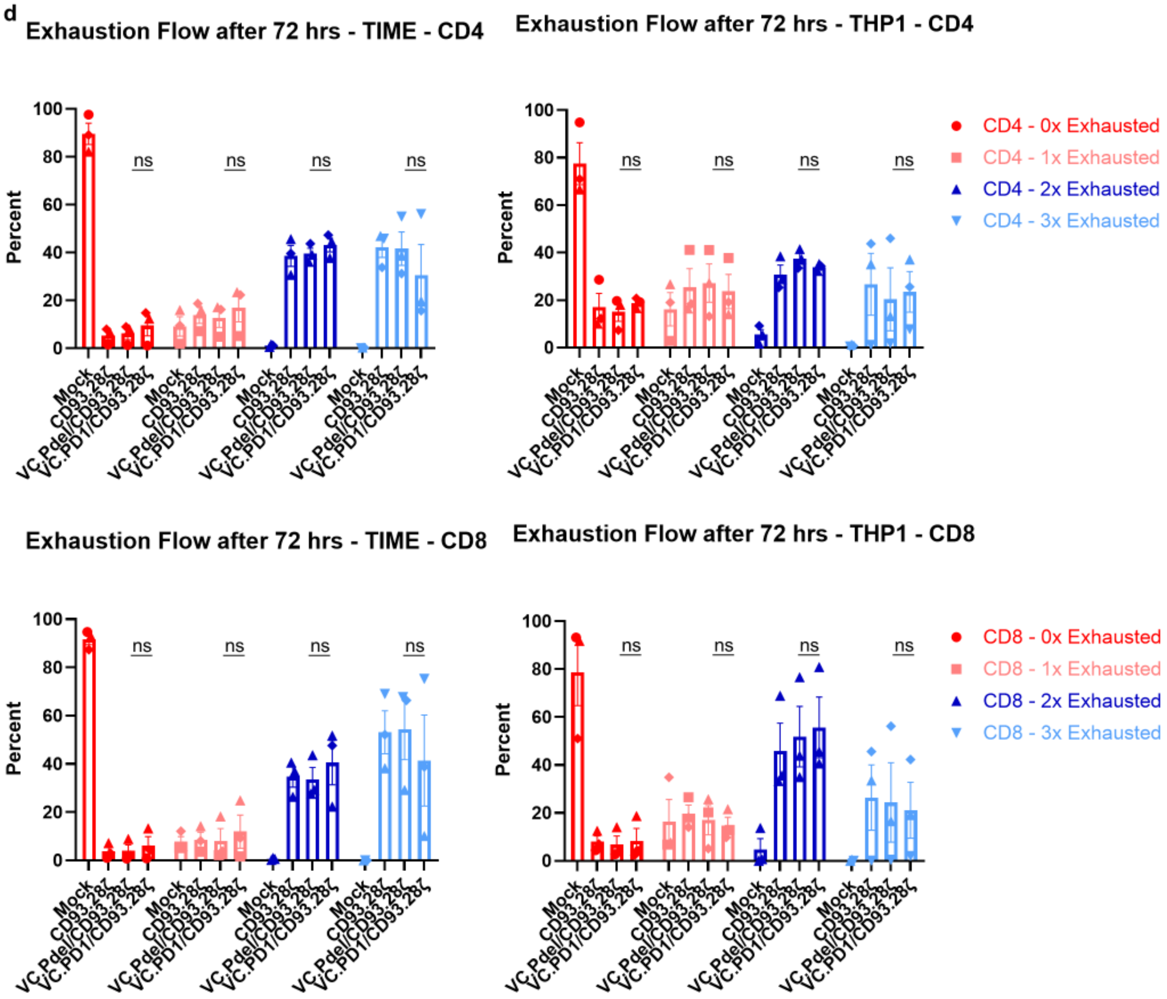
Exhaustion Flow after 72 hrs - TIME - CD4
Exhaustion Flow after 72 hrs - THP1 - CD4
Percent
CD4 - 0x Exhausted
CD4 - 1x Exhausted
CD4 - 2x Exhausted
CD4 - 3x Exhausted
Exhaustion Flow after 72 hrs - TIME - CD8
Exhaustion Flow after 72 hrs - THP1 - CD8
CD8 - 0x Exhausted
CD8 - 1x Exhausted
CD8 - 2x Exhausted
CD8 - 3x Exhausted

e

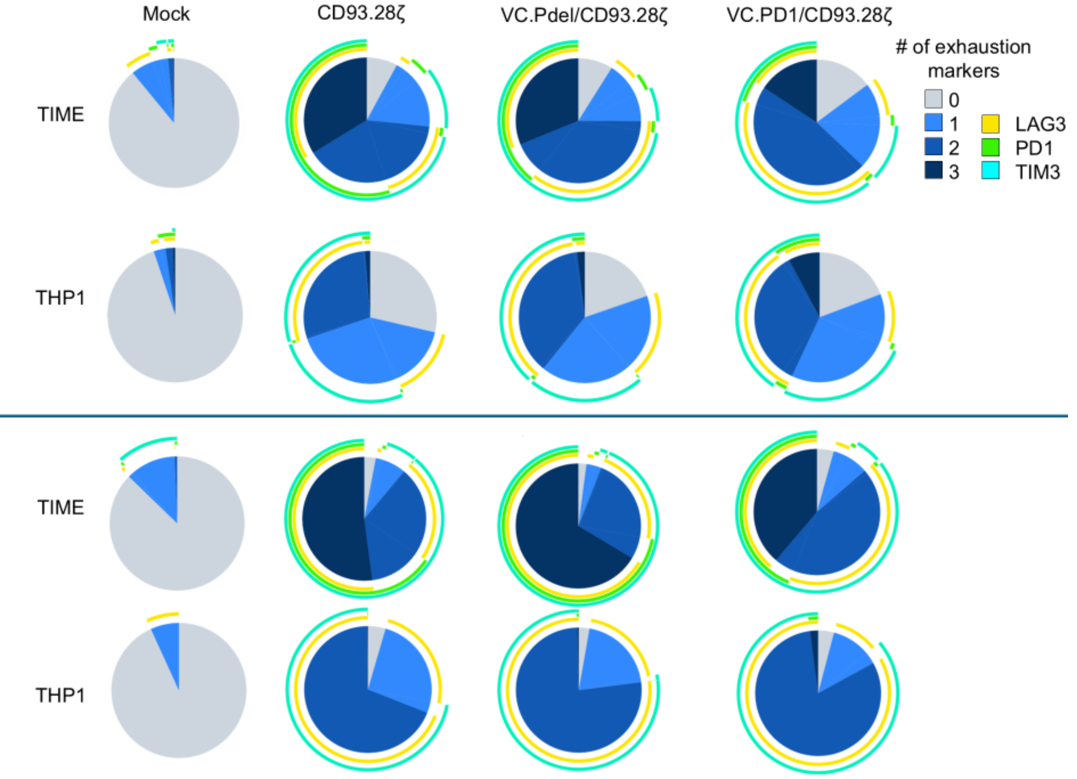


f

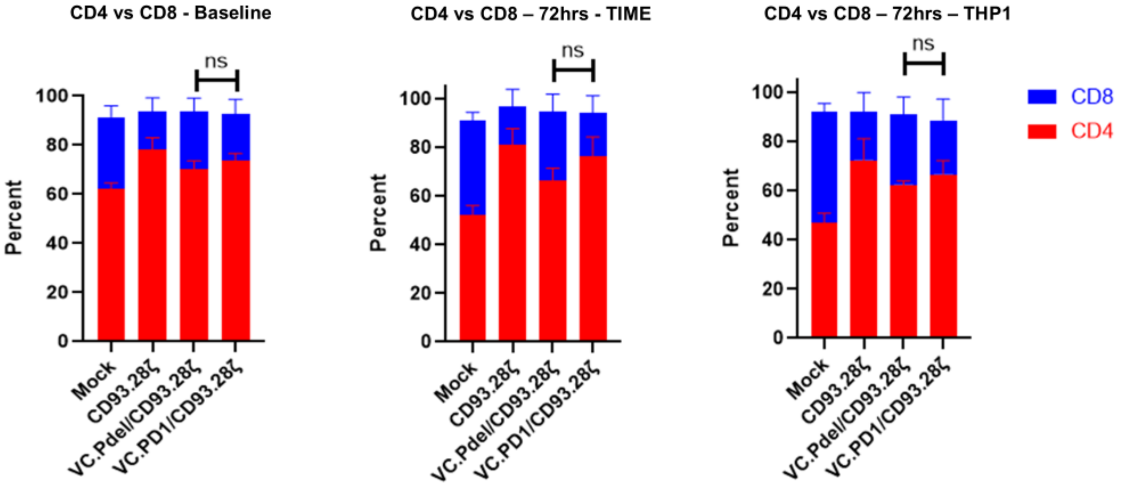


**Supplement 5. A,** Cumulative expression of T cell exhaustion markers (PD1, TIM-3 and LAG-3) measured by flow cytometry after 72 hr co-culture with either TIME or THP-1 cells at 1:2 E:T

ratio. Summary data of n=3 donors is shown. Individual donors are shown with different shapes. No significant difference in exhaustion of VC.PD1/CD93.28ζ and VC.Pdel/CD93.28ζ was observed (unpaired T-test). **B,** SPICE plots of T cell exhaustion markers (PD1, TIM-3 and LAG-3) measured by flow cytometry after 72 hr co-culture with either TIME or THP-1 cells at 1:2 E:T ratio {Roederer, 2011 #275}. One representative donor is shown. **C,** Expression of individual T cell exhaustion markers (PD1, TIM-3 and LAG-3) measured by flow cytometry after 72 hr co-culture with either TIME or THP-1 cells at 1:2 E:T ratio, separated into CD4+ and CD8+ subpopulations. Summary data of n=3 donors is shown. Individual donors are shown with different shapes. No significant difference in exhaustion of VC.PD1/CD93.28ζand VC.Pdel/CD93.28ζ was observed (unpaired T-test). **D,** Cumulative expression of T cell exhaustion markers (PD1, TIM-3 and LAG-3) shown in Supp Fig. 5D, separated into CD4+ and CD8+ subpopulations. Summary data of n=3 donors is shown. Individual donors are shown with different shapes. No significant difference in exhaustion of VC.PD1/CD93.28ζand VC.Pdel/CD93.28ζ was observed (unpaired T-test). **E,** SPICE plots of T cell exhaustion markers (PD1, TIM-3 and LAG-3) measured by flow cytometry after 72 hr co-culture with either TIME or THP-1 cells at 1:2 E:T ratio, separated into CD4+ and CD8+ subpopulations {Roederer, 2011 #275}. One representative donor is shown. **F,** Ratio of CD4+ and CD8+ T cells was measured by flow cytometry at baseline as well as after 72 hr co-culture with either TIME or THP1 cells at 1:2 E:T ratio. No significant difference in CD4-CD8 ratio was observed between VC.PD1/CD93.28ζ and VC.Pdel/CD93.28ζ in all conditions (unpaired T-test).

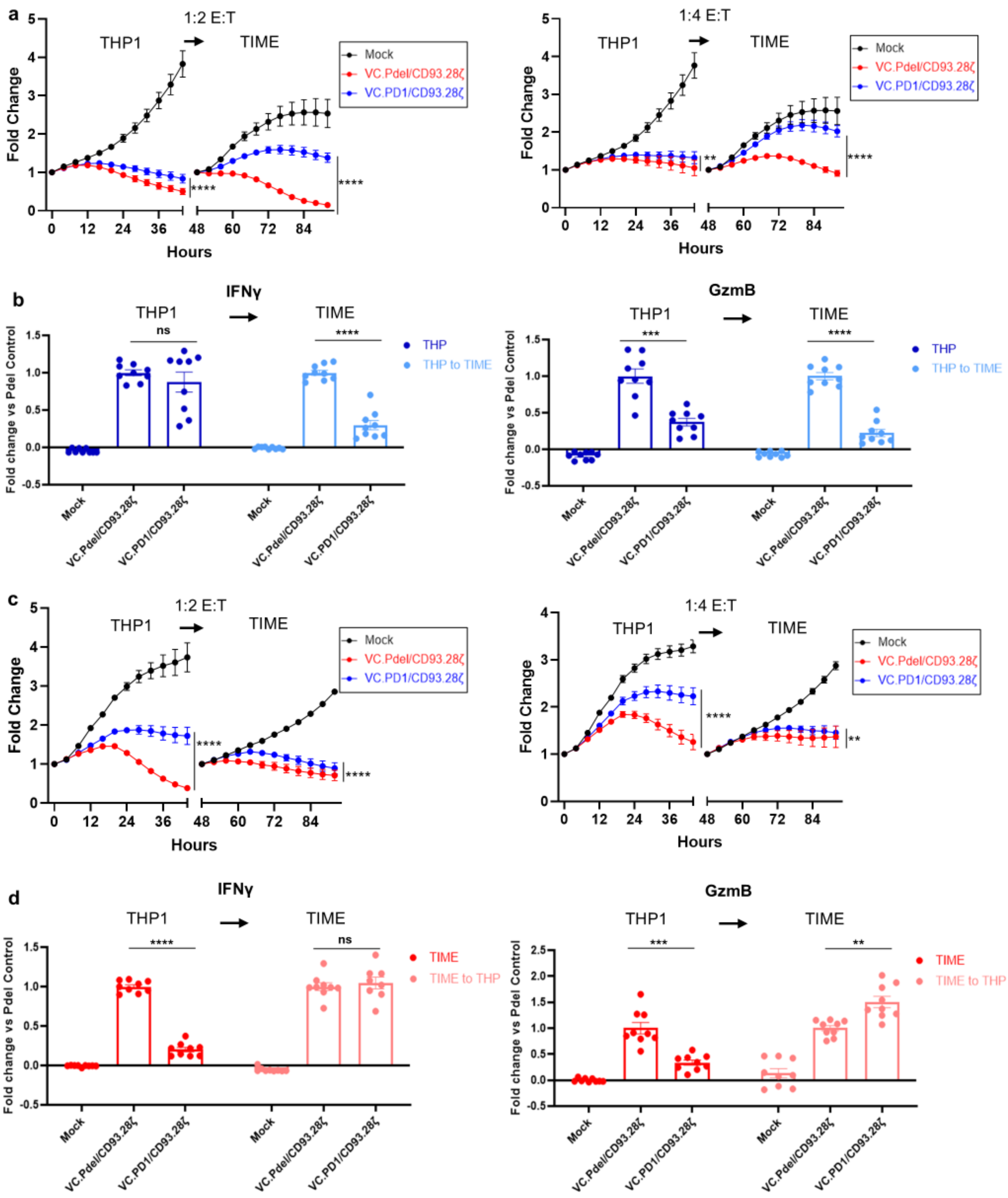


**Supplement 6. A,** Summary Incucyte data from Figure 4A. Data from n= 3 donors at both 1:2 and 1:4 E:T ratios is shown (One-way ANOVA of AUC). **B,** Effector cytokine levels in supernatants taken at 44- and 92-hour timepoints of co-culture with TIME or THP-1 cells at a 1:4 E:T ratio. Cytokine levels depicted as fold change compared to Pdel control, normalized on each assay date (one-way ANOVA). **C,** Summary Incucyte data from Figure 4C. Data from n= 3 donors at both 1:2 and 1:4 E:T ratios is shown (One-way ANOVA of AUC). **D,** Effector cytokine levels in supernatants taken at 44- and 92-hour timepoints of co-culture with THP-1 or TIME cells at a 1:4 E:T ratio. Cytokine levels depicted as fold change compared to Pdel control, normalized on each assay date (one-way ANOVA).
